# Wnt signaling alters CTCF binding patterns and global chromatin structure

**DOI:** 10.1101/2024.01.22.576696

**Authors:** Anna Nordin, Chaitali Chakraborty, Mattias Jonasson, Orgena Dano, Gianluca Zambanini, Pierfrancesco Pagella, Silvia Remeseiro, Claudio Cantù

## Abstract

Wnt signaling plays a pivotal role during development, stem cell maintenance, and tissue homeostasis. Upon Wnt pathway activation, β-catenin translocates to the nucleus where it binds the TCF/LEF transcription factors to drive the context-specific expression of Wnt target genes. Coordinating gene expression programs in vertebrates requires a complex interplay between the regulatory and the 3D organization of the genome. However, the impact of Wnt signaling on genome structure has been poorly explored. Here we investigated how Wnt signaling activation influences the binding patterns of CTCF, one of the core architectural proteins that helps establish the 3D genome organization be demarcating topologically associated domains (TAD). This study uncovered a series of CTCF rearrangements under Wnt, that we termed RUW. Notably, RUW sites that were gained upon Wnt activation were typically dependent on β-catenin and were characterized by both CTCF and TCF/LEF binding. Accordingly, many CTCF RUWs aligned with β-catenin binding patterns, and β-catenin and CTCF co-localized in vivo in discreet nuclear puncta only upon pathway activation. Genome-wide investigation of CTCF-mediated 3D genomic interactions upon Wnt pathway stimulation supported the role of the identified RUWs in mediating Wnt-dependent chromatin loops. Lastly, targeted disruption of selected CTCF binding sites demonstrated their functional contribution to Wnt target gene regulation, implicating regulation of the 3D genomic structure in the execution of transcriptional programs orchestrated by developmental pathways.

## Main

The human body is composed of trillions of cells, all containing an identical genome sequence. The remarkable ability of cells to differentiate into hundreds of specialized cell types is attributed to numerous factors, of which transcriptional regulation is the primary determinant ^1^. Transcriptional regulation therefore dictates cell identity, function, and behavior. This complex process depends on the coordinated actions of signaling molecules, receptors, secondary messengers, and transcription factors that work in tandem to form intricate signaling pathways. Yet, these pathways only represent a fraction of the diverse processes that govern cellular behavior. Technological advances have revealed the role of epigenetic modifications and three-dimensional (3D) chromatin structure in transcriptional regulation, adding another layer of complexity to our understanding of cellular behavior ^2^.

Wnt signaling encompasses three highly conserved signaling pathways that are activated by WNT ligands. Canonical Wnt signaling, referred to as the β-catenin dependent pathway, plays a crucial role in cell communication, modulating various cellular responses like proliferation, migration, and differentiation ^3,4^. During development, Wnt/β-catenin signaling directs early cell fate determination and gastrulation, later driving processes like axis patterning and organogenesis ^5^. In adults, Wnt signaling maintains tissue homeostasis by regulating adult stem cells ^6,7^. Germline mutations that aberrantly activate the Wnt/β-catenin pathway can lead to developmental disorders, while somatic mutations can contribute to the development of cancer ^8^. Colorectal cancer is a well-known example of a Wnt/β-catenin-driven cancer, with over 80% of cases exhibiting loss of function in the tumor suppressor gene *APC* ^9^. At the molecular level, the Wnt/β-catenin signaling pathway is inactive in the absence of WNT ligands. The β-catenin destruction complex, which incorporates APC and the kinase GSK3, phosphorylates β-catenin, causing its ubiquitination and subsequent degradation. This results in low levels of free β-catenin. Upon binding of WNT ligands to cell surface receptors, the destruction complex is recruited to the cell membrane, inactivating it. β-catenin then builds up in the cytosol and translocates to the nucleus, where it binds to the TCF/LEF family of four transcription factors and acts as a transcriptional co-activator ^10^. Although numerous Wnt/β-catenin target genes are well-known and seemingly ubiquitous, accumulating evidence is indicating that the nuclear response of Wnt signaling is largely tissue-specific ^11^. The underlying mechanisms responsible for this specificity are not well understood, representing a crucial area of research for enhancing our comprehension of cell biology and developing therapeutic interventions for Wnt/β-catenin-driven diseases.

The architecture of the genome plays a crucial role in gene regulation. How DNA is packed into chromatin and organized in 3D domains determines the accessibility of DNA for transcriptional complexes, and thus influence the transcriptional rate of genes ^12^. During interphase, each chromosome occupies its own territory, which is further organized into active and inactive compartments, as well as topologically associating domains (TADs). TADs are three-dimensional structures that organize the genome into functional and physical units by bringing together genes and regulatory elements that interact primarily with each other ^13,14^. TADs are thought to form through loop extrusion: as cohesion moves along the chromatin fiber, it traps a loop of chromatin within its ring-like structure until a pair of CCCTC binding factor (CTCF) molecules is reached, which dimerize to anchor and stabilize the loop ^15–17^. Although TADs are large in size, similar processes also occur at a smaller and more transient scale during chromatin loop formation, often between enhancers and promoters or between regulatory elements ^18^. The binding profiles of CTCF have been extensively studied in various organisms and cell types, and despite differences among species, many CTCF binding sites are highly conserved, indicating their important functional roles ^19,20^. While enhancer-promoter looping can occur through other mechanisms, studies have shown that CTCF along with cohesins can either block or facilitate such interactions, with long-range interactions being more dependent on CTCF than short-range interactions ^21^. Even small changes in CTCF binding patterns can affect local gene regulation during stem cell differentiation ^22^, EMT ^23^, and in cancer ^24,25^. Although many of these contexts are connected to Wnt signaling, the specific role of CTCF in Wnt signaling, in particular concerning CTCF’s ability to regulate the 3D genome, is not understood.

Here we performed CUT&RUN ^26^ for CTCF upon activation of Wnt signaling in HEK293T cells and discovered sets of <u>R</u>earrangements of CTCF binding sites <u>U</u>nder <u>W</u>nt signaling (RUW). RUW showed dependence on the presence of β-catenin and enrichment for both CTCF and TCF/LEF motifs, highlighting them as true Wnt-driven events. Accordingly, these regions aligned well with chromatin regions that present increased accessibility to the Tn5 transposase upon Wnt activation ^27^ and with β-catenin and LEF1 binding patterns ^28^. Consistently, physical proximity of CTCF and β-catenin was identified only upon Wnt signaling activation. Next, we performed CTCF HiChIP and not only confirmed the connection between RUW and novel chromatin loops occurring upon Wnt activation, but also revealed larger-scale chromatin structural changes that prompt a need for future research and exploration in other contexts and models. Lastly, we tested the necessity of CTCF binding at RUWs located in the enhancers of the classical Wnt targets *AXIN2* and *DKK1*. Using CRISPR/Cas9 to disrupt the CTCF binding sites reduced the transcriptional response to Wnt activation, proving their regulatory function.

## Results

### CUT&RUN reveals changes in CTCF binding profiles upon Wnt signaling activation

In our previous efforts to chart the genome-wide binding profile of β-catenin, we noticed enrichment for CTCF binding motifs within the β-catenin peak regions ^29^. This finding drove us to hypothesize that Wnt signaling could use CTCF-mediated chromatin organization to regulate target gene expression. To investigate this, we performed CUT&RUN in HEK293T cells to map CTCF genome-wide binding behavior under two conditions: either upon chemical inhibition of PORCN by LGK (Wnt-OFF) or of GSK3 by the small molecule CHIR99021 (Wnt-ON) (N = 3 per condition) (Fig. 1A). After DNA fragment isolation, we performed size selection to enrich for fragments < 120 bp, which represent the footprint of the transcription factor binding directly to DNA ^30^, and identified regions of significant read enrichment using the CUT&RUN peak calling software SEACR ^31^. The concordance of the CTCF replicates in each condition was high (> 80%), consistent with the known stability of CTCF binding. We employed a simple majority approach (2 of 3 datasets) to call peaks: ∼20,000 peaks were identified in each condition, corroborating existing literature ^20^. Of these, over 17,000 (ca. 85%) were present in both Wnt-OFF and Wnt-ON conditions (Fig. 1B). Within the shared peaks, *de novo* motif search with MEME-ChIP ^32^ revealed motifs matching CTCF with an E-value of 4.5 e-4129 (Fig. 1D, top), validating the precision of our datasets.

**Figure 1.**
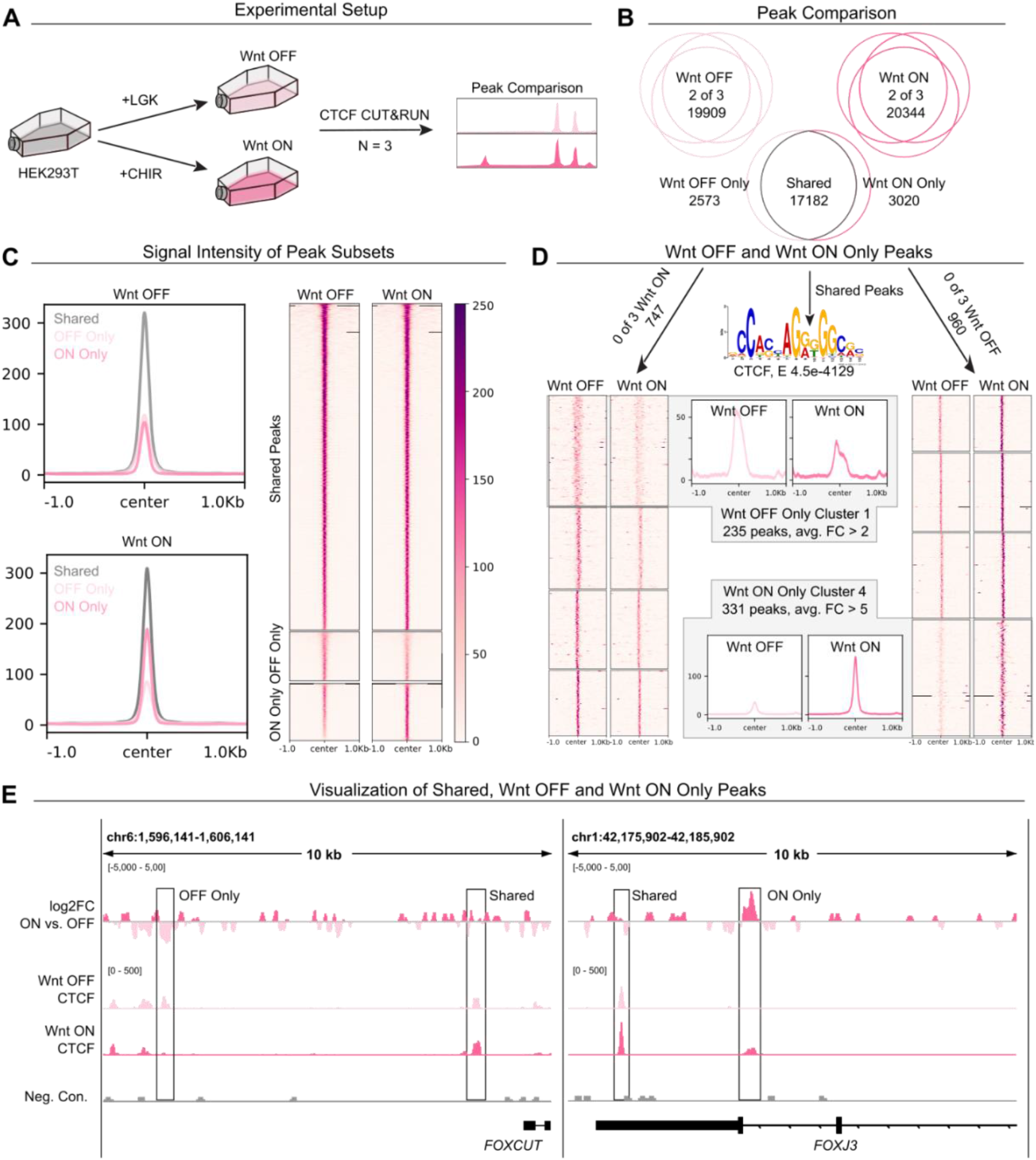
CUT&RUN of CTCF binding in Wnt-OFF vs. Wnt-ON. A. Schematic depicting the experimental strategy. LGK was used to induce Wnt-OFF, and CHIR99021 was used to activate Wnt-ON. 3 independent biological replicates were performed per condition. B. CTCF peak comparison of Wnt-OFF and Wnt-ON. The 2 out of 3 approach was used to select peaks, of which 17,182 were shared while 2,573 were unique to Wnt-OFF and 3020 unique to Wnt-ON. C. Peak average profiles (left) and signal intensity plots (right) showing CTCF signal within shared, Wnt-ON only, and Wnt-OFF only peaks. D. Increased stringency of identification followed by k-means clustering revealed 2 clusters with the highest differences between signal in Wnt-OFF and Wnt-ON. E. Visualization of genomic loci containing Wnt-OFF only, shared, and Wnt-ON only peaks.

We then turned our attention to the CTCF binding events occurring in Wnt-OFF and Wnt-ON. Based on this initial peak calling we identified 2,573 and 3,020 unique peaks, respectively (Fig. 1B). However, most of these peaks presented residual signal in the other group: several regions displayed higher but not exclusive signal to Wnt-OFF or Wnt-ON, indicating that most of these peaks could result from differential binding affinity across the two conditions, rather than *de novo* binding sites in either of them. Notably, the Wnt-ON dataset showed higher average signal intensity in its unique regions (Fig. 1C), suggesting that Wnt signaling induction generates increased affinity of CTCF to a specific set of genomic regions. To identify Wnt-OFF and Wnt-ON exclusive *de novo* peaks, we rendered our selection more stringent by first excluding peaks that were called in a single dataset of the opposite condition. This resulted in 747 peaks unique to Wnt-OFF and 960 in Wnt-ON. Next, we performed k-means clustering (k = 4 clusters) to group these unique peaks based on difference in signal (Fig. 1D). This revealed 1 cluster for each condition with the highest average fold-change (FC) between the two conditions. Cluster 1 in Wnt-OFF with 235 peaks had an average FC > 2 over Wnt-ON, while Cluster 4 in Wnt-ON with 331 peaks had an average of FC > 5 over Wnt-OFF (Fig. 1D). Visual inspection of these peaks confirmed that the nearby shared peaks are comparable between the Wnt-OFF and Wnt-ON, while our approach isolated instances where the unique peaks are convincingly different in their signal profiles (Fig. 1E).

### β-catenin dependent CTCF rearrangements under Wnt (RUW)

Our chosen method of Wnt signaling induction via stimulation with CHIR99021 (CHIR) is known to activate the pathway via inhibition of GSK3 and thus stabilization of β-catenin; however as GSK3 inhibition also stabilizes other proteins and is involved in other pathways ^33,34^, we decided to perform CTCF CUT&RUN in HEK293T cells lacking β-catenin (Δβ-catenin, from Doumpas et al. 2019) to discriminate β-catenin dependent and independent events (Fig. 2A).

**Figure 2.**
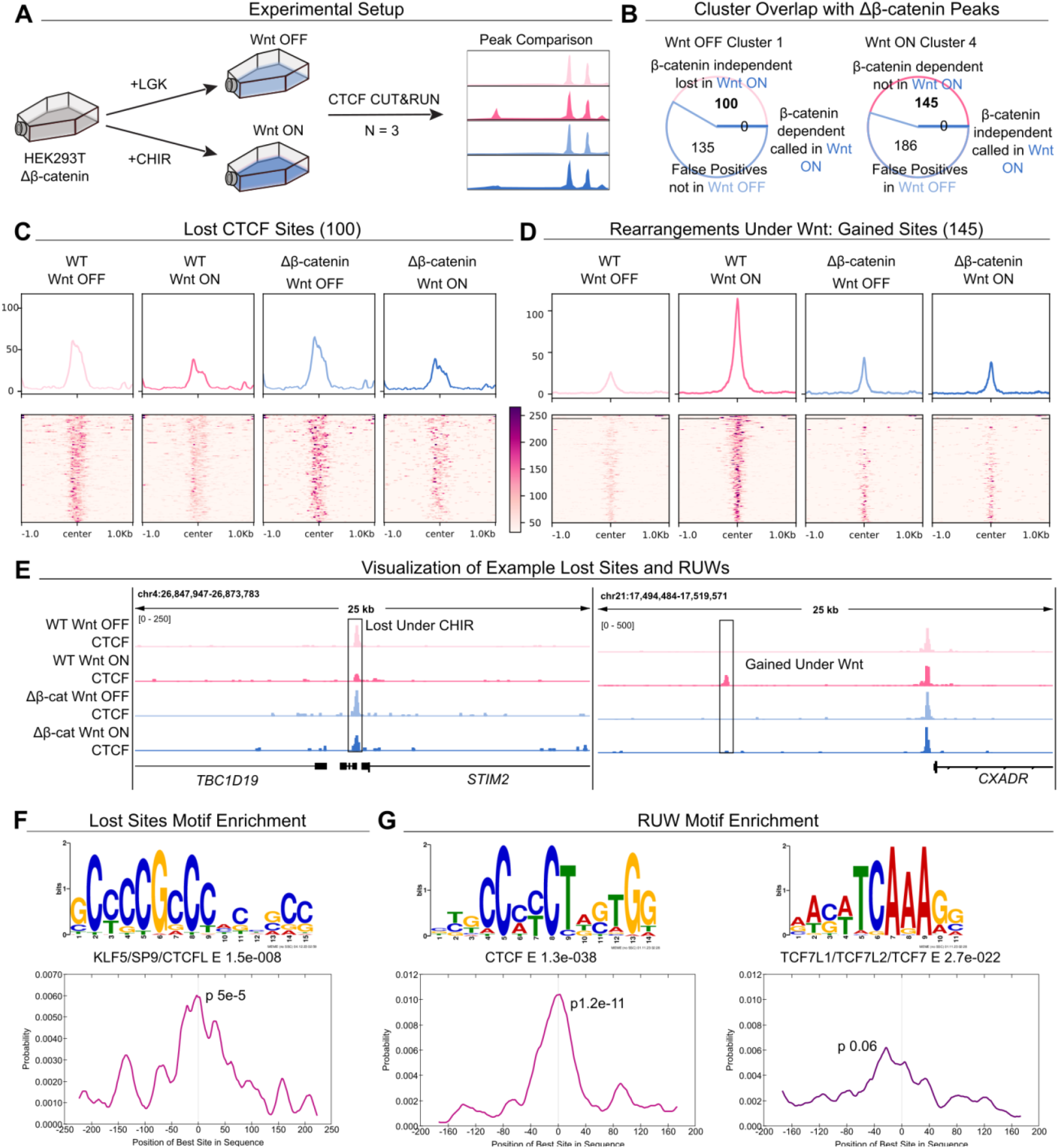
Definition of Rearrangements Under Wnt (RUW). **A.** Schematic depicting the experimental strategy. **B.** CTCF peak comparison of Cluster 1 and Cluster 4 peaks with Δβ-catenin CTCF peaks, defining 100 CTCF sites lost in a β-catenin independent manner, and 145 CTCF sites gained in a β-catenin dependent manner (RUWs). **C.** Peak average profiles (top) and signal intensity plots (bottom) showing CTCF signal within lost sites. **D.** Peak average profiles (top) and signal intensity plots (bottom) showing CTCF signal within gained sites (RUWs). **E.** Visualization of genomic loci containing a lost CTCF site (left, STIM2 locus) and a RUW (right, CXADR locus). **F.** De novo enriched motif (top) within lost CTCF sites matching KLF5/SP9/CTCFL, and centrality within peaks (bottom). **G.** De novo enriched motifs within gained RUWs matching CTCF (left) and TCF/LEF (right) and their centrality enrichment within the peaks (bottom).

We focused on our previously identified Cluster 1 (Wnt-OFF only) and Cluster 4 (Wnt-ON only) sets of peaks, and tested whether they were detected in the absence of β-catenin. Considering Cluster 1, we reasoned that Wnt-OFF CTCF peaks should also be present in Δβ-catenin cells in Wnt-OFF condition, where β-catenin, by definition, does not play a role. Peaks that did not meet this criterion were considered false positives and removed. This left us with 100 peaks, present in Wnt-OFF, but absent in Wnt-ON. Notably, none of these peaks were found in Δβ-catenin cells in Wnt-ON, indicating that the loss of CTCF binding upon CHIR treatment was independent of β-catenin (Fig. 2B, Supp. Table 1), and could be due to other pathways. When considering Cluster 4, we reasoned that if a peak was a true Wnt-ON only phenomenon, it should *not* be present in Wnt-OFF Δβ-catenin: those that were present were also considered false positives and removed. This left 145 CTCF sites that are gained upon CHIR treatment. Of these, none were present in Wnt-ON in Δβ-catenin cells, indicating that their appearance was dependent on β-catenin (Fig. 2B, Supp. Table 2), and thereby a true Wnt-induced event. We termed these 145 sites as CTCF Rearrangements Under Wnt (RUW).

Signal profile and intensity plots, which measure signal enrichment in regions whether a peak is called or not, confirmed the patterns indicated by the peak calling. CTCF binding sites lost under CHIR stimulation showed decreased CTCF occupancy regardless of β-catenin presence, whereas gained sites only showed increased signal in Wnt-ON when β-catenin was present (Fig. 2C, 2D). When normalized and visualized via the Integrative Genome Viewer (IGV, Robinson et al. 2023), nearby CTCF sites are comparable between cell lines and conditions while the lost or gained show differences in signal (Fig. 2E). Next, we performed motif analysis on the lost and gained CTCF binding sites. Lost sites, which are β-catenin-independent, showed enrichment for one *de novo* motif (E 1.5e^-8^), which could be matched with significant p-values to many known motifs, including those of the transcription factors KLF5, SP9, and CTCFL, and showed central enrichment within the peak regions (p 5e^-5^) (Fig. 2F). RUW gained sites had two significantly enriched motifs: one matching CTCF (E 1.3e^-38^) which was centrally enriched (p 1.2e^-11^), and one matching TCF/LEF (E 2.7e^-22^) which was most enriched slightly offset (−40 bp) of peak center (p 0.06) (Fig. 2G). The finding that TCF/LEF motifs frequently lied adjacent to those of CTCF in a close but not overlapping manner indicated that both factors could simultaneously occupy the same region and strengthened the evidence of these CTCF RUWs being Wnt-driven occurrences.

### RUW sites overlap with characterized Wnt responsive regions

We set out to explore the characteristics of Wnt-driven, β-catenin-dependent CTCF RUWs, given their potential relevance in the transduction of Wnt-target genes. To this aim, we first annotated them based on their genomic positions. The majority of RUWs were located in introns, followed by intergenic and promoter regions (Fig. 3A, top). Next, we explored RUW regions based on their chromatin characteristics by measuring how they were marked by the histone modifications H3K4me3, H3K4me1, and H3K27ac with CUT&RUN LoV-U (Fig. 3A, center). As expected, H3K4me3 signal was mostly found in promoter RUWs, which increased in signal upon Wnt induction, indicating that these promoters become more active. H3K4me1 enrichment within RUWs was generally low, though the signal within promoters seemed to decrease upon Wnt signaling activation. As H3K4me1 is typically depleted from active promoters, we considered it consistent with the increase in H3K4me3 ^36^. Many RUWs were decorated with H3K27ac, a marker of open chromatin ^37^: these increased upon Wnt/β-catenin induction (Fig. 3A). We recently did an extensive time course study where we mapped the chromatin state via ATAC-seq upon Wnt/β-catenin signaling ^27^, mapping open chromatin ^38^. We cross-referenced the RUW sites with their chromatin accessibility data, comparing the number of RUWs called as ATAC-seq peaks and the ATAC-seq signal within RUW peaks (Fig. 3B, left). We saw that while most promoters were already accessible and did not change their signal profiles after 24 hr of Wnt induction, some intron and intergenic RUWs showed gained chromatin accessibility (Fig. 3B, left). In combination, these data suggested that the RUW sites possess functional activity.

**Figure 3.**
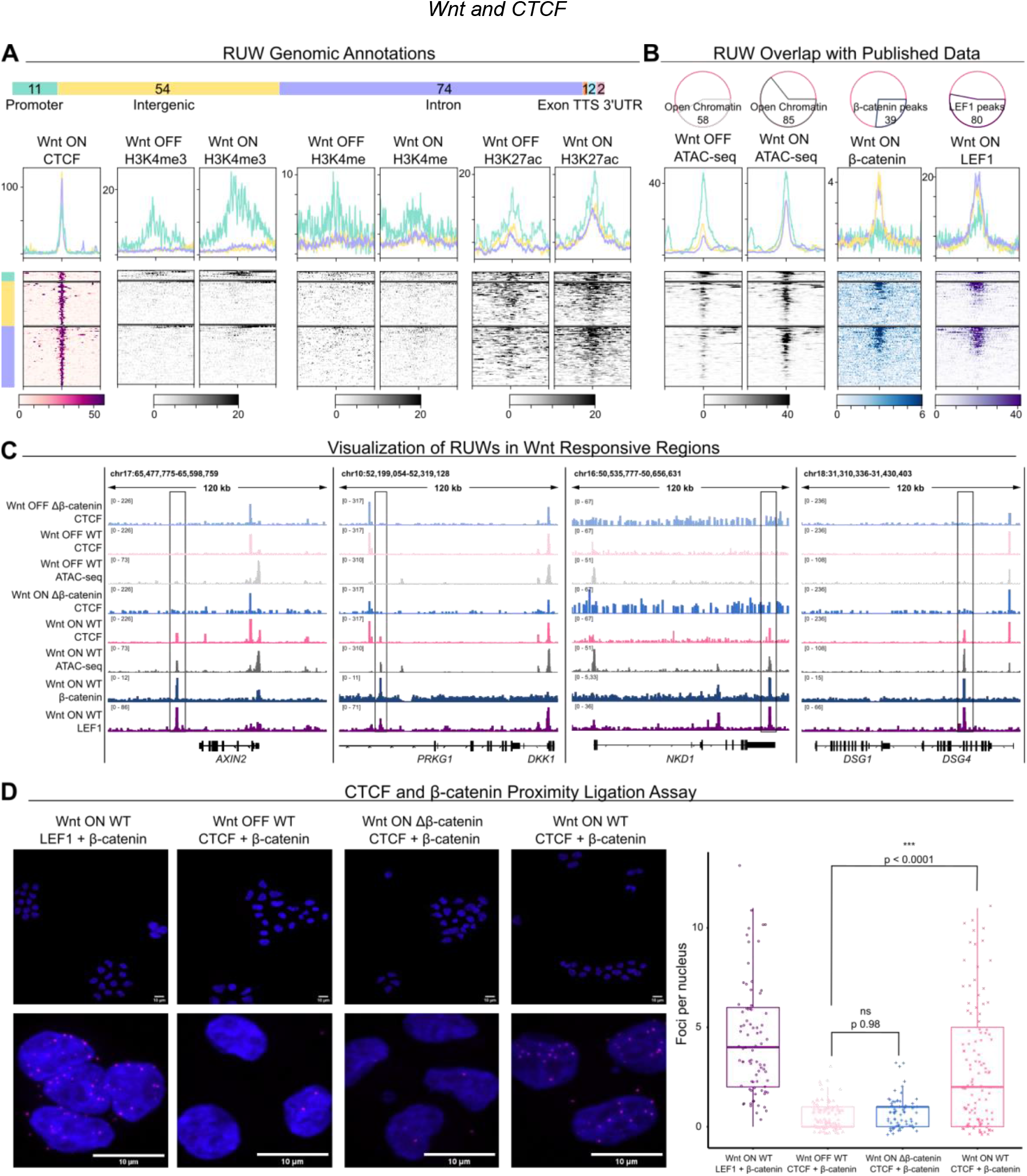
Characterization of RUW regions. **A.** Top: Annotation of RUWs based on genomic location. Bottom: Signal profiles and intensity plots of CTCF, H3K4me3, H3K4me1 and H3K27ac within subsets of RUW peaks. **B.** Left: Venn diagrams of RUW sites called as ATAC-seq peaks, and signal profiles and intensity plots of ATAC-seq signal within RUW peaks. Right: Venn diagrams of RUW sites called as β-catenin and LEF1 peaks, and signal profiles and intensity plots of β-catenin and LEF1 signal within RUW peaks. **C.** IGV tracks of RUW sites that show Tn5 accessible chromatin only in Wnt-ON, β-catenin, and LEF1 binding. **D.** Left: Representative microscopy images from the proximity ligation assay, showing LEF1 + β-catenin in Wnt-ON, and CTCF and β-catenin in Wnt-OFF, Wnt-ON Δβ-catenin, and Wnt-ON WT. Scale bars 10 μM. Right: Quantification of PLA signal, in foci per nucleus. Wnt-ON WT had significantly more interactions than Wnt-OFF or Wnt-ON Δβ-catenin. Counted nuclei = 354.

### CTCF and β-catenin come into physical proximity upon Wnt activation

A comparison of β-catenin, LEF1 and CTCF genome-wide binding profiles revealed – as the motifs indicated – that RUW sites could also be bound by components of the Wnt/β-catenin nuclear complex: > 25% of them were called as β-catenin peaks, and > 50% as LEF1 peaks (Fig. 3B, right). The strongest signal for β-catenin and LEF1 was seen in intron and intergenic RUWs (Fig. 3B, right). RUW sites that were found to exhibit changed chromatin accessibility, β-catenin binding, and LEF1 binding included the notable Wnt targets *AXIN2, DKK1* and *NKD1*, as well as previously unreported direct targets such as *DSG4* (Fig. 3C). The overlapping genomic signal between CTCF and β-catenin/LEF1 suggested a physical interplay between the Wnt transcriptional complex and CTCF. However, while CUT&RUN identifies regions bound by these different factors, it could not distinguish if they ever co-occupy the same locus at the same time. To test this, we performed the highly sensitive proximity ligation assay, which uses microscopy to detect a signal emerging when two proteins of interest are in close physical proximity (within 40 nm) ^39^. Indeed, we could see that β-catenin and CTCF were detected as proximal, and only upon Wnt signaling induction (p < 0.0001, Fig. 3D).

### RUW associated genes include differentially expressed classical Wnt target genes

To further investigate the potential impact of RUWs, we used GREAT ^40^ to assign the RUW peaks to 254 genes (Fig. 4A, Supp. Table 3). STRING mapping revealed a significant degree of interaction within the network (PPI 2.33e^-15^) and Wnt pathway components were overrepresented (FDR 0.0082), making up the center most interconnected cluster (Fig. 4A). Gene ontology analysis also revealed enrichment for the Wnt pathway, and terms related to Wnt signaling such as limb development or cell migration (Fig. 4B). To explore the potential effect of RUWs on gene expression, we overlapped the genes associated to the RUW peaks with gene expression data from HEK293T in Wnt-ON vs. Wnt-OFF (DEG, Log2FC > 0.6, adj. p < 0.05; Doumpas and colleagues ^29^). Of the 254 RUW genes, 20 were DEGs (3.1-fold greater than expected by chance: hypergeometric test, p 9.09e^-6^), consisting of both up- and down-regulated genes (9 up, 11 down), including many of the targets with the highest log2FC such as *AXIN2, DKK1* and *NKD1* (Fig. 4C). To test whether the expression of RUW associated genes was dependent on β-catenin and/or TCF/LEF, we analyzed the DEGs upon Wnt induction in Δβ-catenin and Δ4TCF cells ^29^. Interestingly, while most of the upregulated targets were dependent on both β-catenin and TCF/LEF, the downregulated targets seemed to be mostly independent (Fig. 4D), suggesting that GSK3-inhibition-dependent downregulation might occur via different mechanisms.

**Figure 4.**
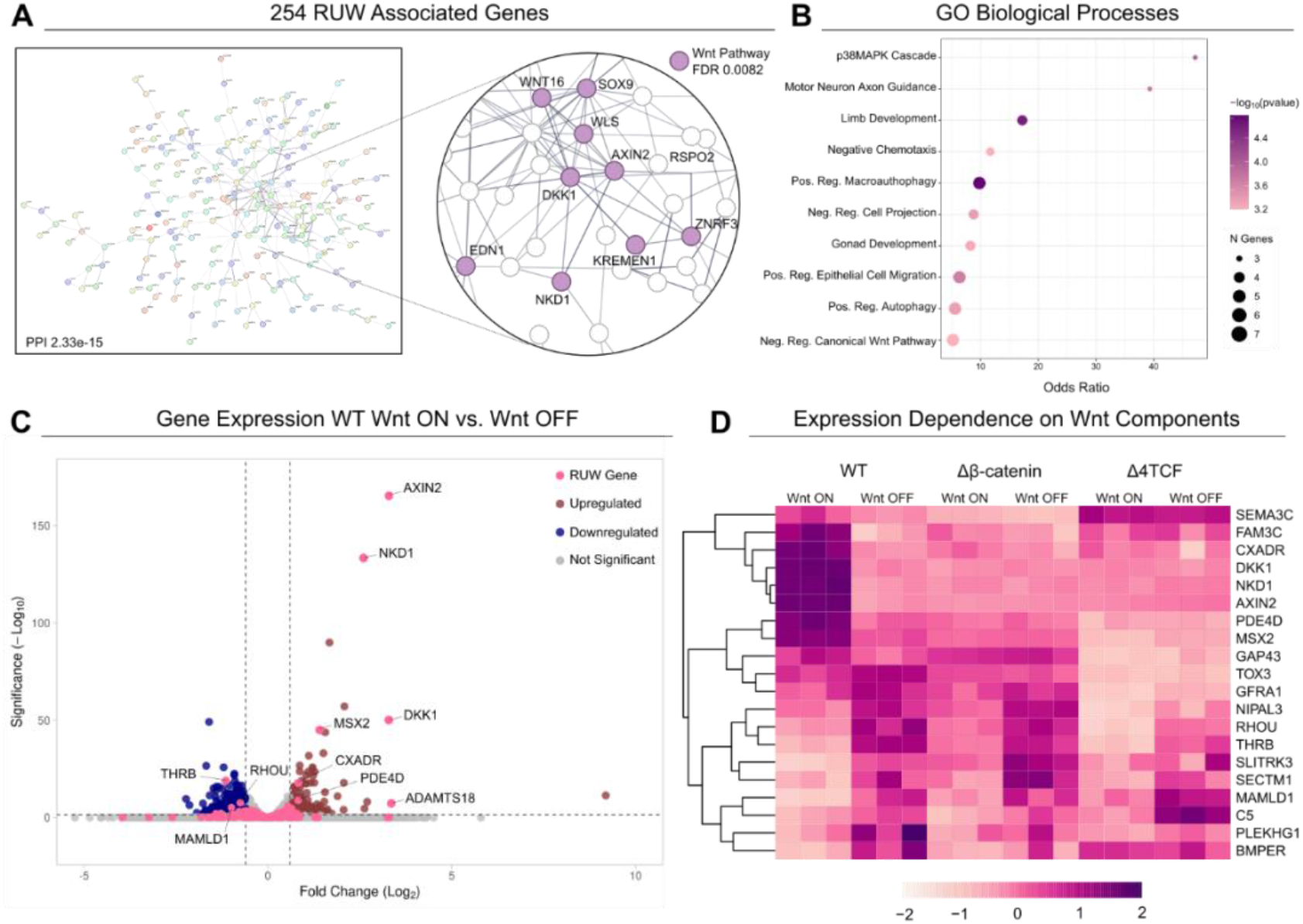
RUW association with genes and gene expression. **A.** Left: STRING map of RUW associated genes. Right: zoom-in on Wnt pathway cluster. **B.** Gene ontology enrichment of RUW genes. **C.** Volcano plot of differentially expressed genes (Log2FC > 0.6, adj. p < 0.05) in Wnt-ON vs. Wnt-OFF. **D.** Heatmap of expression of RUW DEGs in WT, Δβ-catenin and Δ4TCF cells.

### Wnt signaling activation leads to larger-scale CTCF-mediated genomic reorganization

CTCF is a known regulator of the 3D genome structure, both in formation of TADs and smaller scale loops between enhancers and promoters ^18^. We set out to determine whether RUWs were associated with changes in the CTCF-mediated 3D structure by performing HiChIP for CTCF in Wnt-OFF and Wnt-ON conditions (N = 2 per condition) and then comparing loops exclusive to Wnt-ON with RUW peak regions (Fig. 5A). HiChIP genome-wide analysis provided a relatively low number of CTCF mediated loops that were common between the Wnt-OFF and Wnt-ON conditions. When all detected interactions were considered, the ones that were shared consisted of ∼ 20% of loops in each condition, while, at high stringency, only ∼ 1.5% of loops were shared (differential loops with FitHiChIP, FDR 0.01) (Fig. 5B). The difference between conditions could be partially due to the dynamicity of the CTCF looping events leading to a low concordance across replicates ^41^, a similar phenomenon of high CTCF ChIP overlap paralleled by a lower CTCF HiChIP concordance between cell types has been reported previously ^42^. However, the broad difference between Wnt-OFF and Wnt-ON is in line with the previously reported large-scale genomic rearrangements upon modulation of GSK3 activity ^43^. Supporting the latter explanation, we have also noticed that the shared loops, unaltered by GSK3 inhibition, were significantly shorter in length compared to the loops unique to either condition (Supp. Fig. 1A), though the loop annotations were similar in distribution across conditions (Supp. Fig. 1B).

**Figure 5.**
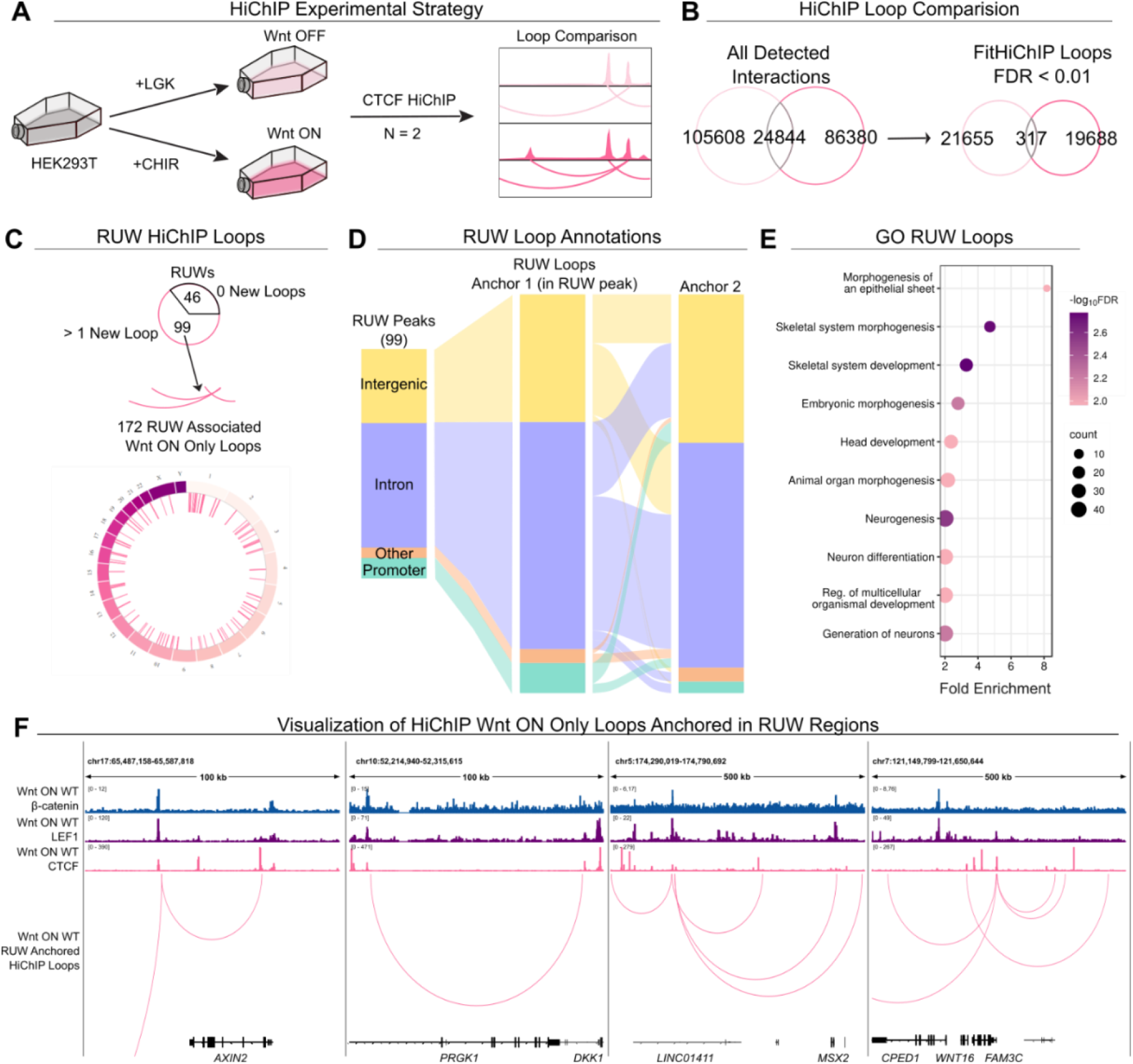
CTCF-mediated 3D genome structural changes upon Wnt activation. **A.** Experimental strategy of performing HiChIP for CTCF in Wnt-OFF and Wnt-ON conditions (N = 2). **B.** Venn diagrams depicting genome-wide comparison of WNT-OFF and WNT-ON HiChIP data. At FDR 0.01 with FitHiChIP, only ∼1.5% of loops in each condition were shared. At lower stringency, this was ∼20%. **C.** Top: Pie chart comparison of RUW peaks with Wnt-ON only HiChIP loop anchors. 99 RUWs were connected to 172 Wnt-ON only loops. Bottom: Circos plot showing genomic locations of RUW associated chromatin loops. **D.** Alluvial plot showing RUW peak annotations, their respective contribution to the RUW loops, and annotations of their connecting anchor. **E.** Gene ontology biological processes of the set of RUW loop anchors, showing enrichment for developmental processes and morphogenesis. **F.** Visualization of RUW associated Wnt-ON only CTCF HiChIP loops at the AXIN2, DKK1, MSX2 and WNT16 loci. None of these loops are present in Wnt-OFF.

We focused our attention on those changes that coincide with our previously stringently identified RUWs, where a CTCF repositioning occurs in Wnt-ON and is dependent on β-catenin. Of the 145 gained RUWs, 99 were connected to other genomic regions via a total of 172 CTCF-mediated loops that are also specific to the Wnt-ON condition (Fig. 5C, Supp. Table 4). The RUWs involved in looping were mostly found within introns and intergenic regions, likely representing enhancers. These RUWs were often an anchor of multiple loops, which were most likely to have the other anchor in intergenic and intron regions, though we identified some instances of enhancer-promoter looping (Fig. 5D). Gene ontology analysis performed on the set of RUW loops revealed that these are likely involved in the coordination of biological processes, including epithelial morphogenesis, neurogenesis and embryogenesis (Fig. 5E), that are all broadly consistent with most Wnt/β-catenin signaling output *in vivo* ^3^.

The RUWs connected to other regions via chromatin loops present only in Wnt-ON included those found within the *AXIN2* and *DKK1* loci (Fig. 5F). The HiChIP data revealed, as we hypothesized, that these RUWs were associated with the chromatin looping towards another CTCF site closer to the promoter of the gene (Fig. 5F). Yet, we also identified instances where many new loops were tethered to RUW sites upon Wnt signaling stimulation, likely helping to connect hubs of TCF/LEF and β-catenin bound regions (Fig. 5F, *MSX2* and *WNT16*).

### Perturbation of CTCF binding sites within RUWs affects Wnt target gene upregulation

To test whether CTCF RUWs are functionally implicated in gene expression, we sought to selectively disrupt their CTCF binding motifs. We selected the RUWs annotated to *AXIN2* and *DKK1*, as they are two of the most differentially expressed genes in our analysis and quasi-universal Wnt/β-catenin targets. The *AXIN2* RUW was located within the characterized enhancer region for the gene ^44^, while the *DKK1* RUW was located in putative enhancer region ∼100 kb away. We identified the exact position of the CTCF and TCF/LEF binding site(s) within these RUW regions and, as indicated by the earlier motif analysis, found that the two motifs and corresponding CUT&RUN peak summits were slightly offset. This allowed us to design CRISPR sgRNAs which would lead Cas9 to only disrupt the core of the CTCF binding motif (Fig. 6A). We transfected cells with Cas9 and either the targeting or a scrambled sgRNA control, and then stimulated to obtain Wnt-OFF and ON conditions. To avoid clonal effects, we measured gene expression in populations (N = 6 independent wells per condition) and could see that the sgRNA targeting the *AXIN2* RUW led to a significant decrease in upregulation of *AXIN2* (p 0.0028) but not *DKK1* nor *NKD1* (Fig. 6B, top panel). Similarly, the sgRNA against the *DKK1* RUW decreased upregulation of only *DKK1* (p 0.02) (Fig. 6B, bottom panel). We confirmed disruption of CTCF binding sites in the populations: the detected mutations included indels and sequence alterations which invariably affected CTCF but not TCF/LEF consensus sequences (Fig. 6C). These results indicate that CTCF binding to RUW regions contributes to the upregulation of each associated Wnt target gene.

**Figure 6.**
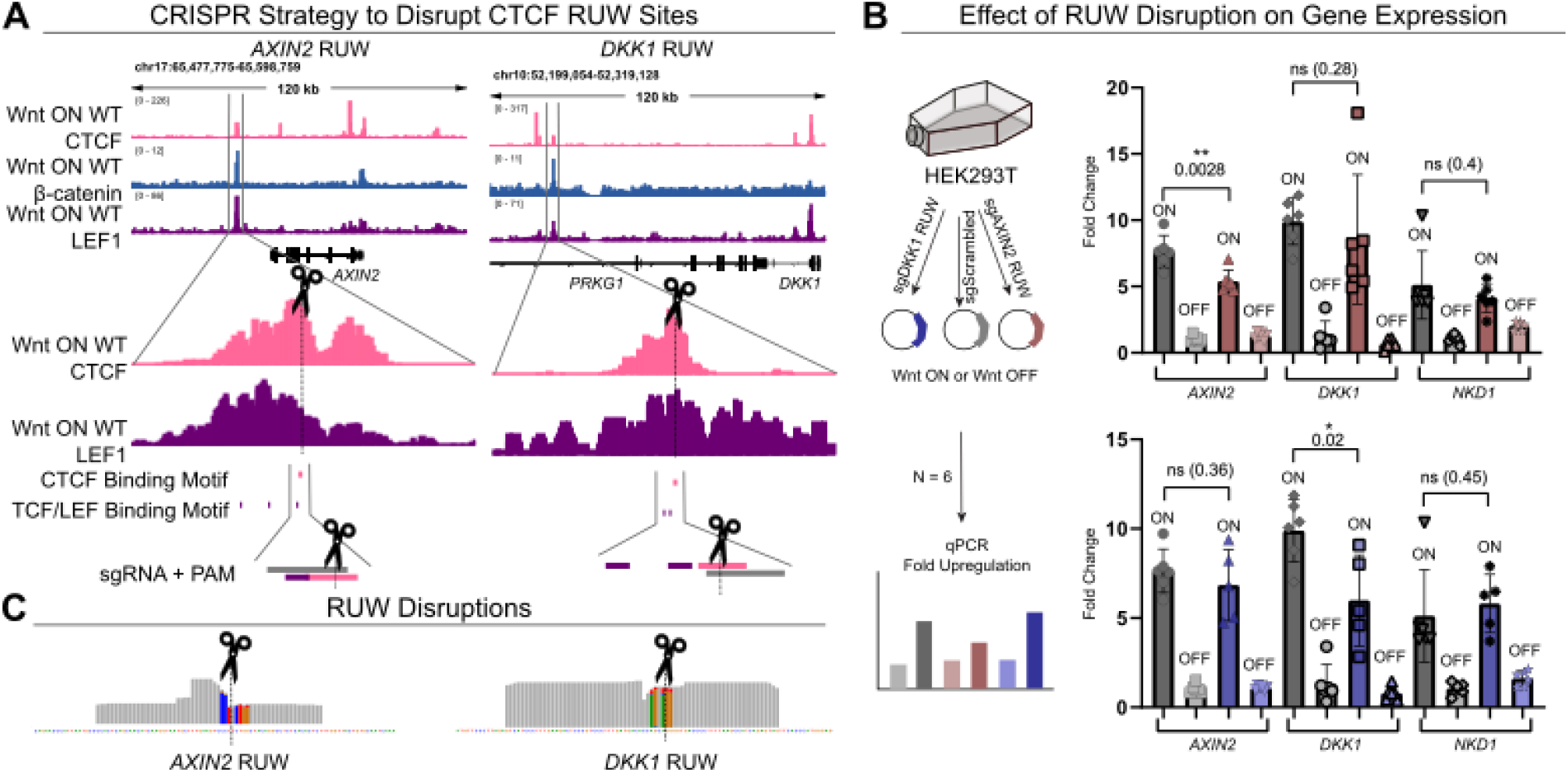
CRISPR/Cas9 disruption of CTCF RUW binding sites. **A.** CRISPR/Cas9 strategy to mutate RUW CTCF binding sites near AXIN2 and DKK1. sgRNAs were designed to disrupt the CTCF motif. **B.** Gene expression measurement by qPCR showing that RUW disruption significantly and only affects the upregulation of the corresponding gene for AXIN2 (p 0.0028) and DKK1 (p 0.02) compared to a scrambled control. **C.** Bam coverage of RUW sites from sequenced populations with confirmed disruptions in CTCF sites (alteration in PAM sequence + 5bp upstream). Basepairs with > 2% of non-matching alleles are colored by the representation of each nucleotide.

## Discussion

Enormous efforts have been dedicated to unraveling the complexity of the response to Wnt/β-catenin signaling activation and deciphering the mechanisms that drive its remarkable tissue-specificity ^45^. This included the identification of the different components of the Wnt/β-catenin transcriptional complex ^46–49^ and recently the varying patterns of β-catenin genome-wide binding in a time and context specific manner ^27,50–53^. However, the impact of Wnt/β-catenin activation on the 3D genome structure and its potential role in modulating the Wnt-dependent transcriptional output, is still poorly understood. Our establishment of CTCF genome-wide binding patterns across Wnt pathway stimulation and the discovery of the 3D genomic interactions that involve CTCF biding sites, lead us a model where, upon Wnt activation, CTCF is detected at newly accessible Wnt responsive cis-regulatory elements. Here, CTCF binds in tandem with β-catenin and TCF/LEF to induce the formation of chromatin loops that contribute to the upregulation of Wnt target genes (Fig. 7). This, to our knowledge, represents a previously unknown mechanism of Wnt target gene regulation.

**Figure 7.**
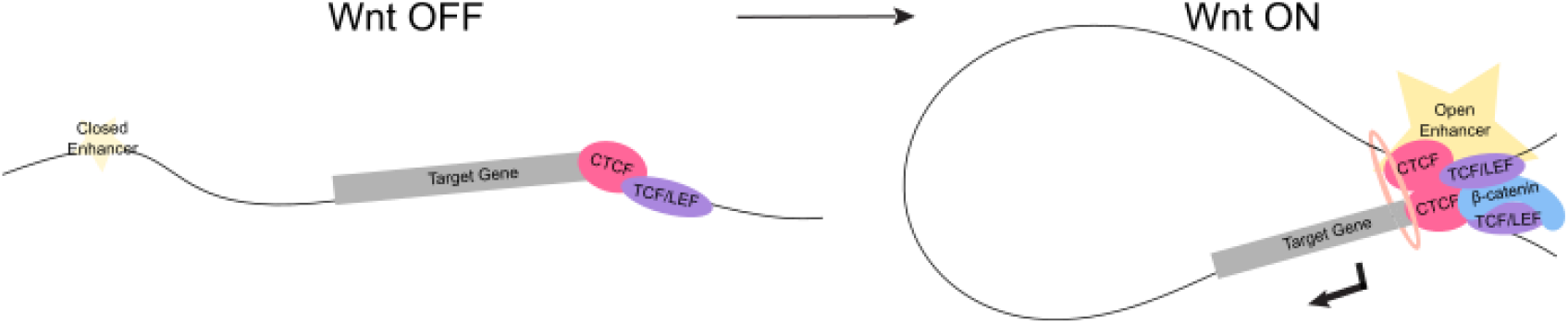
Rearrangements Under Wnt (RUW) model. Left: When Wnt is OFF, the chromatin is closed at enhancers of Wnt target genes and transcription is low. CTCF binding is constitutive at a region near the promoter of the gene. Right: When Wnt is activated, the chromatin at the enhancer opens and TCF/LEF and β-catenin bind. CTCF binds the RUW site and mediates enhancer-promoter looping, which boosts target gene expression.

While the process by which CTCF is recruited to these sites remains to be solved, our data obtained in Δβ-catenin cells, where CTCF does not bind RUW sites upon GSK3 inhibition, indicates that the physical presence of β-catenin is necessary for CTCF rearrangements. This would imply that β-catenin, possibly together with other components of the Wnt transcriptional complex, acts as pioneer to open the chromatin and recruit CTCF, and not vice versa. While a full-time course of CTCF binding upon Wnt activation would be needed to determine the time dynamics of RUWs, β-catenin binding to these sites early (90 minutes or 4 hours; Pagella et al. 2023) supports the model in which β-catenin reaches these sites before CTCF. It has previously been reported in the literature that CTCF can bind to or interact with β-catenin ^25^, TCF7 ^54^, and the member of the ChiLS complex LDB1 ^55^, all of which could thus be considered as potential partners in recruiting CTCF to the chromatin.

Recent characterization of the *AXIN2* enhancer – which also corresponds to one of our identified RUWs – has been performed ^44^. This study included the introduction of mutations throughout the whole enhancer sequence followed by testing the activity of the mutated variants in a luciferase-dependent Wnt transcriptional reporter. The core of the CTCF binding site that we have identified is located between 288 - 292 bp into the enhancer region characterized by the Cadigan group (see figure 5A in Ramakrishnan et al. ^44^): mutating this region in their study showed no change in the ability of the enhancer to activate the reporter. Consistently, the *in vitro* luciferase reporter, while it successfully detected enhancer activity, took the enhancer out of its 3D context of the genome such that one would expect *not* to find a notable contribution of the 3D chromatin structure to transcriptional activation. Our results would therefore imply that only in its native context does this enhancer – and presumably all the other in correspondence to the identified RUWs – need the CTCF binding site to sustain full regulatory activity ability. This lends credibility to the idea that CTCF, at least in the context of Wnt target gene regulation, is acting as a mediator of genome organization and not as a classical transcription factor ^20^.

Interestingly, it has previously been reported that CTCF mediated chromatin loops are instructive in the formation and maintenance of liquid-liquid phase separated condensates ^56^ and also that β-catenin becomes a component of these nuclear condensates upon Wnt activation at enhancers of its target genes ^57^. This raises the possibility that the RUW-mediated chromatin loops could play a role in the formation of phase-separated transcriptional condensates at Wnt target genes, which might be responsible for driving transcription, an interesting theory to be explored in future research.

In this study, we focused on RUWs that align with the *AXIN2, DKK1, NKD1* genomic loci, as they likely represent the most reliable ubiquitous Wnt target genes (https://web.stanford.edu/group/nusselab/cgi-bin/wnt/target_genes_components). However, as Wnt target genes appear to depend on context ^51,53^ and time ^27^, only by exploring other systems and models of Wnt activation can we uncover whether distinct sets of RUWs exist, thereby deciphering whether involvement of CTCF constitutes a mechanism for Wnt signaling to exert its tissue-specific responses. Alternatively, as the genes we have considered, *AXIN2, DKK1* and *NKD1*, all encode for Wnt pathway negative feedback regulators, it is also possible that CTCF will emerge as a general, and perhaps ubiquitous, facilitator of the negative feedback regulation of the pathway itself.

The number of high-confidence β-catenin-dependent RUWs we identified was relatively small when compared to the initial ∼3000 CTCF CUT&RUN peaks present only in Wnt-ON condition. As the Wnt-ON condition was achieved by CHIR administration, many of these 3000 events are independent of β-catenin, pointing to a potentially larger effect of GSK3 inhibition on the 3D genome structure than the direct effect of canonical Wnt signaling. This is also consistent with our genome-wide analysis carried out by CTCF-HiChIP that identified a vast number of CTCF-dependent chromatin loops that follow treatment with CHIR. The proposition that WNT ligands might act *in vivo* as selective inhibitors of GSK3, and that stabilization of β-catenin is only one of the several consequences triggered ^58–60^ would imply that broad CTCF-mediated 3D genome organization changes will be observed in many other *in vitro* and *in vivo* setups. Our study therefore underscores the need to investigate the role of CTCF and of the 3D genome organization that follows the response to Wnt activation across all cellular contexts in which this pathway plays a key role.

## Acknowledgments

The authors are grateful to Stefan Koch and Giulia Pizzolato for the help and reagents used to conduct the PLA assay. This work was supported by Grants to Cl.Ca. from Cancerfonden (CAN 2018/542 and 21 1572 Pj), the Swedish Research Council, Vetenskapsrådet (2021–03075 and 2023-01898), Linköping University and the Knut och Alice Wallenbergs Stiftelse. This project was fostered by collaborative grants to S.R. and Cl.Ca. from the National Molecular Medicine Fellows Program (NMMP, now PALS, for Program for Academic Leaders in Life Science). S.R. and Cl.Ca. are Fellows of the Wallenberg Centers for Molecular Medicine (WCMM) and receive generous financial support from the Knut and Alice Wallenberg Foundation. The computations and data handling were enabled by resources provided by the National Supercomputer Centre (NSC), funded by Linköping University. Peter Münger at the National Supercomputer Centre is acknowledged for assistance concerning technical and implementational aspects in making the codes run on the Sigma resource.

## Author contributions

A.N., M.J. and Cl.Ca. conceived the project. A.N., M.J., Ch.Ch and O.D. performed the experiments. A.N. performed the formal analyses, prepared the figures and drafted the first version of the article. Ch.Ch. conducted and analyzed HiChIP. G.Z., P.P. and S.R. provided critical scientific and methodological input. S.R. supervised the HiChIP experiments and provided financial support for the study. Cl.Ca. supervised the research team and provided financial support for the study. All authors reviewed and commented on the final manuscript.

## Competing interest statement

The authors declare no competing interests.

## Methods

### Cell Culture

HEK293T human embryonic kidney cells (parental) and Δβ-catenin cells (generated in Doumpas et al. 29) were cultured in a 37 °C incubator in 5% CO2 and 89% humidity. Culture medium used was high glucose Dulbecco’s Modified Eagle Medium (Cat. #41965039, Gibco) supplemented with 10 % bovine calf serum (Cat. #1233C, Sigma-Aldrich) and 1X Penicillin-Streptomycin (Cat. #15140148, Gibco). Cells were stimulated either with 10 μM CHIR99021 (Wnt-ON) or 1 nM LGK (Wnt-OFF) for 24 hr prior to harvest for assays.

### CUT&RUN

CUT&RUN for CTCF was performed as described in 26, with minor modifications as described here. Three independent rounds of experiments were performed for the three biological replicates of each cell line and condition. 250,000 cells were used per replicate. Antibodies used included anti-CTCF (abcam, ab70303) and anti-HA (Merck, 05-902R) at 1:100 dilutions. For each replicate of CTCF, a corresponding anti-HA negative control was performed from cells in the same well. 40 μl of ConA beads were used per sample, and digitonin concentration was 0.025% throughout the protocol. After fragment release, DNA was purified with phenol:chloroform:isoamyl alcohol followed by ethanol precipitation, and stored until library preparation.

CUT&RUN LoV-U for H3K4me3, H3K4me1 and H3K27ac was performed according to Zambanini et al. 28 based on the original protocol from the Henikoff lab 26, with minor modifications as described here. Experiments were performed in duplicate for each condition. 250,000 cells were used per sample and bound to 10 μl magnetic ConA agarose beads. Antibodies used included anti-H3K4me3 (antibodies online, ABIN6971977), anti-H3K4me1 (antibodies online, ABIN3023251), and anti-H3K27ac (antibodies online, ABIN2668475) at 1:100 dilutions. The digestion buffer was kept and added to the release fraction before DNA purification. DNA purification of H3K27ac samples was done with phenol:chloroform:isoamyl alcohol followed by ethanol precipitation.

Library preparation was performed according to Zambanini et al. 28 using the KAPA Hyperprep Kit (KAPA Biosystems, Cat. #KK8504) with the following modifications: for CTCF samples, 15 cycles of amplification were performed, while LoV-U samples received 13 cycles as previously described. For CTCF samples, size selection was performed post-amplification using 2% E-Gel Size Select II Gels (Thermo Fisher, Cat #G661012), retrieving fragments from 150 – 240 bp (representing 30 – 120 bp without the adapter sequences). Samples were sequenced with 36 bp pair-end reads on the NextSeq 550 (Illumina) using the Illumina NextSeq 500/550 High Output Kit v2.5 (75 cycles) (Cat. #20024906, Illumina). Library preparation and sequencing of the CTCF samples was all done in parallel with sequencing on the same lane.

### CUT&RUN Data Analysis

Reads were trimmed with bbmap bbduk (version 38.18, ^61^) to remove adapter sequences, known artifacts, [AT]18, [TA]18, and poly G and C repeats. Alignment to the hg38 genome was performed using bowtie (version 1.0.0, 62) with the options -v 3 -m 1 -X 120 for CTCF and with -X 350 for the histone modification LoV-U samples. SAMtools (version 1.11, 63) was used for bam file creation, mate fixing, and deduplication. Bedgraphs were created with BEDtools (version 2.23.0, 64) genomecov on pair-end mode. Peaks were called using SEACR (version 1.3, 31) on norm relaxed mode against the corresponding negative control. Peak sets were subsequently filtered to remove peaks falling within CUT&RUN suspect list regions 65. Peak overlaps were performed using BEDtools.

For graphs and visualization purposes, replicate bam files were merged with SAMtools into a single file. deepTools (version 3.5.1-0, 66) was used to convert bam files to normalized bigwig files (bamCoverage using -RPGC option), make log2FC difference tracks (bamCompare), signal intensity plots and profiles (computeMatrix followed by plotHeatmap) and to perform clustering (plotHeatmap -kmeans, 4 clusters). Genome region annotation was done with HOMER (version 4.11, 67) on default settings. Motif analyses were performed with the MemeChIP suite 32: Centrimo was run on non-local mode, both the motifs used for meme identification and FIMO results were used to identify specific motif instances within peaks. Peak-gene annotation was performed using GREAT (version 4.0.4, 40) on default settings. STRING 68 with disconnected nodes removed and SR Plot (https://www.bioinformatics.com.cn/srplot) were used for data visualization.

### Published Data Integration

RNA-seq data was downloaded from Doumpas and colleagues 29and processed in R (version 4.2.2) for data visualization. ATAC-seq bigwig data were downloaded from Pagella and colleagues 27 and signal within RUW regions explored using deepTools. CUT&RUN data of LEF1 and β-catenin were downloaded from Zambanini et al. 28; peaks were called with SEACR with a threshold of 0.05 for comparison with RUWs.

### Proximity ligation assay (PLA)

Proximity ligation was performed using Duolink PLA assay kit (Merck, Cat. #DUO92102) according to the manufacturer’s guidelines. Antigens were detected using mouse anti-β-catenin (BD Labs, 610154), rabbit anti-CTCF (abcam, ab70303), and rabbit anti-LEF1 (antibodies online, ABIN1680678) antibodies at 1:200 dilutions. NucBlue (Invitrogen, Cat. #R37606) was used to counterstain for nuclei. LEF1 and β-catenin in WT Wnt-ON was used as a positive control of the assay, while CTCF and β-catenin in WT Wnt-OFF and in Δβ-catenin Wnt-ON were used as negative controls. The assay was performed in duplicate. Images were acquired using the Zeiss LSM 700 confocal microscope, imaging nuclei and PLA foci on separate channels. Acquisition settings were optimized on the positive control and remained the same for all samples. Images were taken with 40X magnification, taking z-stacks of 10 slices of 1 micrometer.

Image processing was performed in ImageJ ^69^ and Fiji ^70^. For figures, z-stack images were combined into a maximum projection, orange was converted to magenta, and scale bars were added. For quantification, channels were split and converted to 16 bit. Thresholding was applied equally to all images. Analyze particles (> 1 micron) was used to count nuclei, while PLA foci were counted using find maxima (> 100), then foci per nucleus were counted using the ROI measure function. Between 67 – 111 nuclei were counted per condition, for a total of 355 nuclei. Statistical testing was done with two-tailed t-tests using GraphPad Prism (version 9.3.0).

### CRISPR/Cas9 RUW Disruption

sgRNAs were cloned into the pX330spCas9-HF1 plasmid. pX330-SpCas9-HF1 was a gift from Yuichiro Miyaoka (Addgene plasmid#108301;http://n2t.net/addgene:108301; RRID:Addgene_108301) Sequences were confirmed via Sanger sequencing. Cells were seeded in 12-well plates 6 hours prior to transfection. Transfections were performed using the calcium phosphate method, with 12 independent transfections performed per sgRNA. 16 hours post-transfection, the cells were washed with PBS and the media was changed. After 24 hours, media was changed to media containing either CHIR99021 or LGK as described above, 6 replicates per sgRNA and condition. After 24 hours, media was removed and cells were lysed with Qiazol (Qiagen, Cat. #79306). RNA extraction was performed according to manufacturer’s guidelines. cDNA conversion was performed using the high-capacity cDNA reverse transcription kit (Thermo Fisher, Cat. #43-668-13) according to manufacturer’s guidelines, converting 1 ug of RNA per replicate. qPCR was performed using the SYBR green PCR master mix (Applied Biosystems, Cat. #4309155) in 10 ul reactions on the BioRad CFX96, all reactions were done in technical duplicate and the Cq values averaged. Relative quantification was performed using the method developed by Pfaffle to account for primer efficiency 71. Housekeeping genes included *GAPDH* and *ACTB*. To calculate fold-change, all relative expression values were compared to the average of the Wnt-OFF scrambled sgRNA values for each gene. Statistical testing was done with two-tailed t-tests using GraphPad Prism. All primer and sgRNA sequences are listed in Table 1.

**Table 1.**
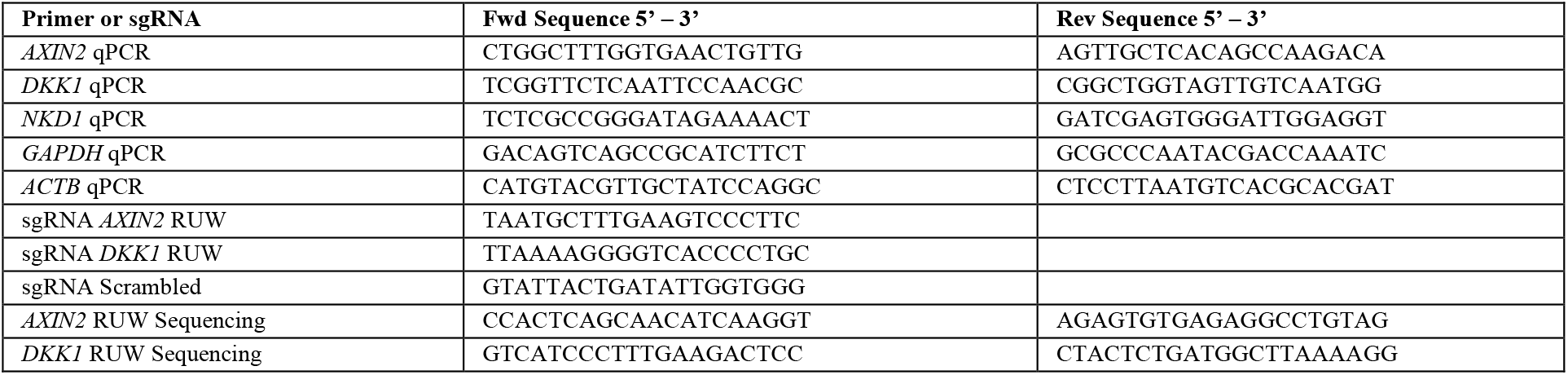
Primer and sgRNA sequences.

### Validation of RUW Disruptions

Primers were designed as to allow for sequencing of the putative Cas9 cut site within the RUW regions with 36 bp reads: the cut site was not located within the primer, but within the following 16 base pairs. For *AXIN2* the cut site was covered by only 1 read, while for DKK1 it was covered by both read directions. Primer sequences are listed in Table 1. RUW regions were amplified from genomic DNA of transfected populations via PCR with the high fidelity Q5 polymerase (New England Biolabs, Cat. #M0491S), with an annealing temperature of 62 degrees and performing 35 cycles. Correct product size and a single product was confirmed with an agarose gel, then the DNA was purified with bead cleanup to remove primers and contaminants. Amplicons were prepared for sequencing as described above for CUT&RUN samples, with the exception that size selection was performed only to remove adapter dimers. Samples were sequenced to ∼1.5 million raw reads.

The Cas9 potential cut site was determined to be within the PAM site plus 5 bp upstream (8 bp total). Reads were trimmed as described above to remove artifacts, and then further trimmed to identify amplicons containing WT sequences (containing the WT 8bp cut site as described above). 71% of *AXIN2* RUW amplicons and 80% of *DKK1* RUW amplicons contained the full WT cut site. The remaining sequences were aligned to the genome, and visualization of the types of mutations and/or indels was done by viewing the bam coverage in IGV, highlighting bp with > 2% alleles with mutations for the figure, confirming that these occur primarily within the cut site (within the CTCF motif) and do not extend further into other regions of the amplicon (such as the TCF/LEF motif of *AXIN2*). Some reads aligned to other regions of the genome or did not align at all, which could underlie off-targets of the PCR or larger indels that resulted in spurious mapping. Due to this and difficulty in quantification of mutations after multiple rounds of PCR (35 cycles of original amplification, 13 cycles during library prep), we did not consider the editing efficiency in the populations could be accurately estimated.

### CTCF HiChIP

HiChIP was performed in duplicates in both LGK (Wnt-off) and CHIR99021 (Wnt-ON) cell lines using the Arima-HiC+ Kit (Arima, Cat. #A101020) and according to the guidelines in the Arima-HiChIP user guide for mammalian cells. HEK293T cells were cultured as described above and stimulated for 24 hours for Wnt-ON and Wnt-OFF, and approximately 5 × 106 cells per line were used to obtain at least 15 μg of input DNA for HiChIP. Cells were fixed in 1% fresh methanol-free formaldehyde for 15 min at room temperature while rotating, and Glycine was used to quench the crosslinking reaction (final concentration 125 mM Glycine) for 5 min. After two washes with cold PBS, crosslinked cells were harvested in PBS by scraping, pelleted and stored at - 80°C until further usage. In the subsequent steps, crosslinked chromatin was digested with restriction enzymes, end-filled with biotinylated nucleotides and ligated. Proximally ligated chromatin was then sheared on a Covaris E22O instrument, with the following shearing parameters: shearing time 5 min, PIP 105, duty factor 5, and 200 cycles per burst, to achieve a fragment size range of 200 bp to 800 bp. Chromatin immunoprecipitation was performed with an antibody against CTCF (AbFlex CTCF, Active Motif, Cat. # 91285, Lot 31321002), using 0.5 mg of antibody per mg of sheared chromatin. After biotin enrichment and adapter ligation, immunoprecipitated DNA was subjected to PCR amplification (18 cycles) using Accel-NGS 2 S Plus DNA Library Kit (#21024, Swift Biosciences) and indexing Kit (#26696, Swift Biosciences), according to the Arima -HiChIP Library Prep user guide. Quality controls for chromatin digestion, ligation and shearing were conducted through gel electrophoresis on a 1.5% agarose gel and library profiles were assessed through Bioanalyzer High Sensitivity DNA Analysis (Agilent #5067–4626). The HiChIP libraries were sequenced on a NovaSeq 6000 Sequencing System (Illumina) aiming ∼400 million 150PE reads per library.

### HiChIP analysis

HiChIP analysis was performed as described before 72,73 using HiC-Pro and FitHiChIP. Fastq files were quality-checked with FastQC (https://www.bioinformatics.babraham.ac.uk/projects/fastqc/, 0.11.8). Adapters and low-quality reads (phred < 33) were removed using cutadapt (v 4.0). Mapping to the human genome (GRCh38/hg38) and retrieval of valid interacting fragments was performed using the HiC-Pro software v3.1.0 and setting ligation sites as GATCGATC, GANTGATC, GANTANTC, GATCANTC, removing duplicates and merging replicates per sample. Valid loops were identified using FitHiChIP v 11.0 and significant loops were filtered at an FDR threshold of 0.01. To identify the differential (Wnt-off/Wnt-ON) and common loops, we ran venn_interactions.r, a utility script from FitHiChIP. The criteria were set such as Wnt-off specific loops are those present only in LGK sample [i.e., ∃ LGK (counts ≥ 1) & ∄ CHIR99021 (counts = 0)], while Wnt-ON specific loops are present only in CHIR99021 sample [i.e., ∃ CHIR99021 (counts ≥ 1) & ∄ LGK (counts = 0)]. Annotation of the significant and differential loops was done using GenomicInteractions package in R (1.26.0) and loops were classified as promoter-promoter (P-P), promoter-enhancer (P-E) or enhancer-enhancer (E-E) setting the criteria for promoter regions <2 kb TSS. Differences in loop length (bp) between differential and common loops were assessed using a Wilcoxon test with Benjamini-Hochberg correction with ggpubr (0.6.0). Among the differential loops, multi-anchor loops were identified as loops that share anchors with other loop(s) i.e., one anchor is utilized by two or more loops. Gene Ontology analysis on identified gene-promoters of differential loops and multi-anchor loops were conducted with clusterProfiler (v 4.10.0).

## Availability of Data and Materials

Raw and processed sequencing data generated in this study have been submitted to ArrayExpress under accession number E-MTAB-13727. Public data used in this study can be downloaded and accessed according to the reference information provided.

## Notes

### Competing Interest Statement

The authors have declared no competing interest.

