## Supplementary Figure 1 for "Wnt signaling alters CTCF binding patterns and global chromatin structure"

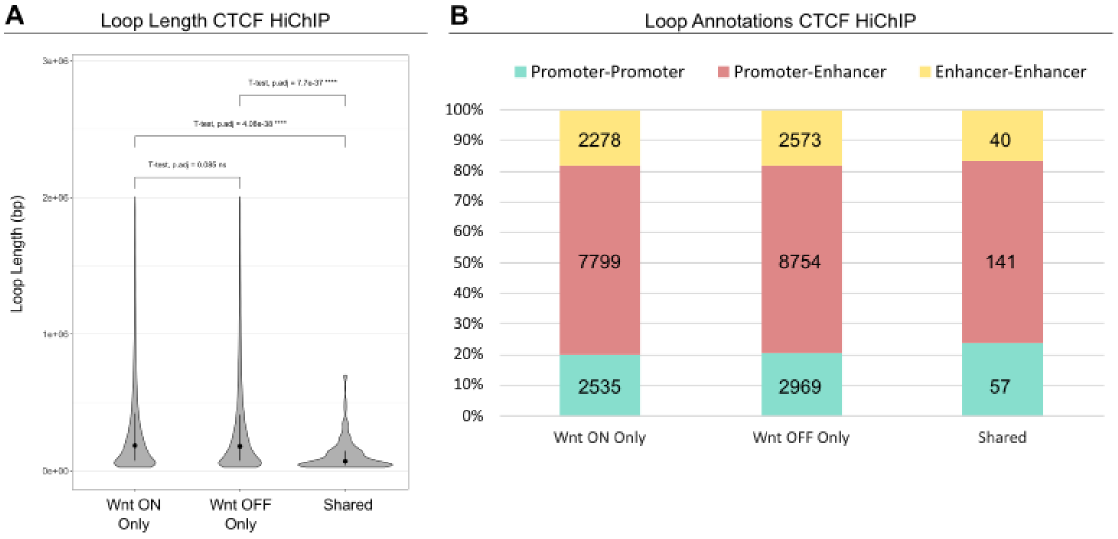


**Supplementary Figure 1. CTCF HiChIP genome-wide analyses. A.** Plot of loop length distribution of Wnt-ON only, Wnt-OFF only and shared loops. Shared loops were significantly shorter than the unique loops. **B.** Loop annotations of CTCF loops across conditions, which were similar in distribution.
